# Enterovirus A71 adaptation to heparan sulfate comes with capsid stability tradeoff

**DOI:** 10.1101/2024.02.23.581741

**Authors:** Han Kang Tee, Gregory Mathez, Valeria Cagno, Aleksandar Antanasijevic, Sophie Clément, Caroline Tapparel

**Author notes:** These authors contributed equally (Aleksandar Antanasijevic, Sophie Clément, Caroline Tapparel).

## Abstract

Because of high mutation rates, viruses constantly adapt to new environments. When propagated in cell lines, certain viruses acquire positively charged amino acids on their surface proteins, enabling them to utilize negatively charged heparan sulfate (HS) as an attachment receptor. In this study, we used enterovirus A71 (EV-A71) as model and demonstrated that unlike the parental MP4 variant, the cell-adapted strong HS-binder MP4-97R/167G does not require acidification for uncoating and releases its genome in the neutral or weakly acidic environment of early endosomes. We experimentally confirmed that this pH-independent entry is not associated with the use of HS as an attachment receptor but rather with compromised capsid stability. We then extended these findings to another HS-dependent strain, suggesting that adaptation to HS generally modifies capsid stability and alters entry mechanism. Our data show EV-A71 pH-independent entry for the first time and, more importantly, highlight the intricate interplay between HS-binding, capsid stability, and viral fitness, wherein enhanced multiplication in cell lines leads to attenuation in hostile *in vivo* environments such as the gastrointestinal tract.

## Introduction

Heparan sulfates (HS) are linear, negatively charged polysaccharides connected to various cell surface and extracellular matrix proteins. Expressed on a wide range of cells, they play a pivotal role in various biological processes, and many viruses exploit them to attach and concentrate onto cell surfaces before binding to main entry receptor^1^. Despite a substantial body of literature on the subject, the actual implication of HS binding on viral infections remains a topic of debate.

Enterovirus-A71 (EV-A71) is an excellent example of the ongoing controversy regarding the impact of HS receptor utilization in viral pathogenesis. This virus is a member of the *Picornaviridae* family and the most neurotropic EV after poliovirus. It causes significant hand, foot and mouth disease outbreaks, particularly in Asia-Pacific countries, and is associated with severe neurological complications, notably in small children and immunosuppressed patients^2^. The virus uses human scavenger receptor class B member 2 (SCARB2) as the main receptor for internalization and uncoating^3,4^. Since SCARB2 is mostly localized on lysosomal membrane and sparsely on plasma membrane^3,5^, it seems to play only a minor role in EV-A71 cell attachment^6^. Consistently, numerous other EV A71 receptors have been described in the literature, including HS^3,7^. When propagated in cell culture, EV-A71 rapidly acquires adaptive mutations (i.e. patches of positively charged amino acids on the viral capsid) that allow them to bind HS, sometimes with high avidity. These strong HS-dependent variants grow efficiently in cell culture but show attenuated virulence in animal models, such as mice and cynomolgus monkeys^8-11^. Analysis of the differential expression of SCARB2 and HS in tissues from monkey or transgenic mice revealed little overlap. Strong HS expression was detected in sinusoidal endothelial cells and vascular endothelia, where SCARB2 was not detected^9,10^. Similarly, HS expression in the brain was mainly found in vascular endothelia but SCARB2 expression was found predominantly in neuronal cells. The authors of these studies concluded that binding to HS on endothelial cells in absence of SCARB2 leads to viral trapping, abortive infection, and attenuation^9,10^. Similar observations were shown for other viruses, including Murray Valley encephalitis^12^, Japanese encephalitis^12^, Sindbis^13^, Theiler’s murine encephalomyeliti^14^, tick-borne encephalitis^15^, West Nile^16^ and dengue^17^.

We previously isolated cell-adapted EV-A71 mutants with strong affinity for HS which emerged upon passaging of intermediate HS binders derived from both patient and mouse-adapted MP4 strains in cell culture^18,19^. The mutants presented two amino acid changes in the VP1 capsid protein: VP1-L97R mutation in the VP1 BC loop, shown to confer intermediate affinity for HS together with a secondary mutation, VP1-E167G, located in the VP1 EF loop, which significantly strengthened HS binding with reduction of negative charges^19,20^. As previously observed for strong HS-binding variants, we showed that, in contrast to the original mouse-adapted MP4 strain which exhibited virulence in mice, this cell-adapted MP4-97R/167G double mutant was completely attenuated in mice^19^. In the current study, we used MP4 and MP4-97R/167G mutant as representatives of respectively, weak and strong HS-binders, slow and fast-growing in cell lines and virulent and avirulent in mouse models (as documented previously^19,20^), to elucidate the consequence of virus adaptation towards HS binding on the viral growth cycle. We demonstrated that these mutations not only increase binding to HS, but also reduce capsid stability, leading to improved uncoating and faster cell internalization in a HS-independent manner. Of note, another strong HS binder harboring VP1-E145Q substitution also showed decreased capsid stability compared to the wildtype HS-independent variant. These data provide another possible explanation for the *in vivo* attenuation of strong HS-binders which may originate from viral trapping but also from decreased capsid stability which would be detrimental to the virus in challenging environments such as the gastrointestinal tract.

## Results

### Lysosomotropic drugs reduce infectivity of the HS-independent MP4 but enhance infectivity of the strong HS-binder MP4-97R/167G

First, we sought to assess whether viruses displaying different dependence on HS exploit different growth cycle pathways. We compared the effect of lysosomotropic drugs, namely hydroxychloroquine (HCQ) and bafilomycin A1 (BAF-A1) on MP4 and MP4-97R/167G double mutant. As presented in **Fig. 1A**, Vero cells were pre-treated for 1 hour with each drug before infection. These drugs showed no cytotoxic effect at the concentrations used in the assay (**Fig. S1A**). Inoculation was then performed for 1 hour in presence of the drug and inoculum was removed and replaced with fresh drug-free media. The number of infected cells was determined at 24 hours post-infection (hpi) using immunofluorescence. To confirm inhibition of endosomal acidification by these drugs, the presence or absence of acidic lysosomes was assessed by immunostaining of lysosomal-associated membrane protein 1 (LAMP1) and by staining with lysotracker, a dye specific for acidic compartments. The amount of LAMP1 and lysotracker double-positive lysosomes decreased drastically following treatment with the drugs, confirming the inhibition of endosomal acidification (**Fig. 1B**). The effect of the drugs on viral replication was then compared for the two variants (**Fig. 1C & D**). MP4 infectivity was significantly reduced by both drugs in a dose-dependent manner, while MP4-97R/167G infectivity was in contrast enhanced. Similar results were obtained in RD cells (**Fig. S1B**), indicating that drug effects are not cell-dependent. The different sensitivity of the two variants to acidification inhibitors was more pronounced with HCQ, so we performed a detailed examination of the mechanism of action with this drug.

**Fig 1.**
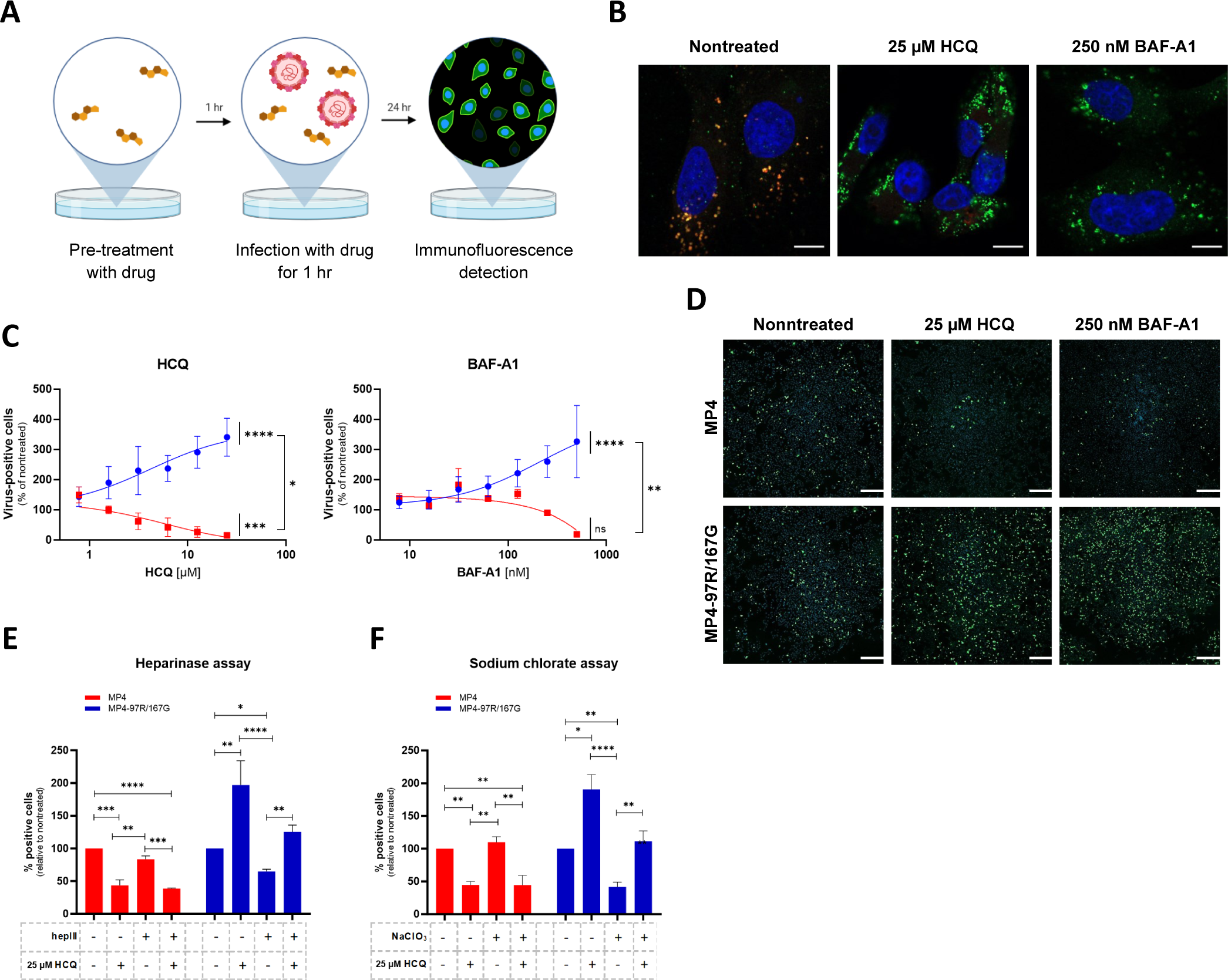
Lysosomotropic drugs inhibit infection by MP4 but not by MP4-97R/167G. (**A**) Schematic illustration of the virus inhibitory assay workflow. Cells were pre-treated with lysosomotropic drugs and infected in presence of the drug. After inoculum removal, infected cells were cultured in drug-free media and infected cells were stained by immunofluorescence (IF) with anti-VP2 Ab. (**B**) Inhibition of endosomal acidification confirmed with lysotracker staining (red). Lysosomes are in green (anti-LAMP1 Ab) and nuclei in blue (DAPI). Representative IF images are shown (scale bar, 10 µm). **C**) Dose response assay in infected Vero cells. Results are shown as % of virus-positive cells relative to nontreated control. Statistical significance (one-way ANOVA) between treated and untreated virus or between treated MP4 and MP4-97R/167G was calculated based on the AUC. (**D**) Representative IF staining of EV-A71 (anti-VP2 in green) 24 hpi of Vero cells in presence of 25 µM HCQ or 250 nM BAF-A1 (scale bar, 300 µm). (**E & F**) HCQ effect in Vero cells pre-treated or not with heparinase III (hepIII) (**E**) or sodium chlorate (NaClO_3_) as in A (**F**). Statistical significance (two-way ANOVA) was calculated for each virus between each condition. In D to F, mean and S.E.M of biological triplicates are shown. *p < 0.05, **p < 0.01, ***p < 0.001, ****p < 0.0001.

To determine whether the effect of HCQ was related to the usage of HS as attachment receptor, we repeated the virus inhibitory assay using cells depleted of HS by either heparinase digestion (**Fig. 1E**) or treatment with sodium chlorate (**Fig. 1F**). The distinct sensitivity to HCQ was reproduced regardless of the presence or absence of HS on the cell surface. Of note, we confirmed that as for the human strain^20^, the HS-independent and HS-dependent MP4- derivatives both need SCARB2 to infect cells and cannot replicate in SCARB2 CRISPR-Cas9 knock-out cells (**Fig. S2**). Altogether, these data indicate that the capsid mutations change the sensitivity to HCQ, independent of the attachment receptor used.

### MP4 enters via SCARB2-mediated and pH-dependent endocytosis, while MP4-97R/167G utilizes an alternative SCARB2-dependent pathway

In our previous studies, we showed that MP4 virus uses SCARB2 as entry receptor and here we demonstrated that it is inhibited by HCQ. This strongly suggests that MP4 uses SCARB2-mediated pH-dependent endocytosis for entry and uncoating as demonstrated for many other EV-A71 variants^7,21^. After having excluded that HCQ effect did not affect virus binding (**Fig. 2A**), we conducted single-cycle infection assay and showed that differential effect of HCQ on MP4 and MP4-97R/167G became prominent from 4 hpi onward (**Fig. 2B**). To further dissect which step of the viral growth cycle was differentially affected by the treatment, we performed a time-of-addition assay. As shown in **Fig. 2C**, HCQ significantly lost its effect when administered later than 2 hpi, confirming that the effect occurs during the early phase of the viral cycle. To more specifically assess whether the drug affects viral entry, we transfected *in vitro* transcribed genomic RNA containing the nanoluciferase (Nluc) gene as reporter (**Fig. 2D**). RNA transfection allows to bypass receptor-mediated entry. In these conditions, no difference was observed for both variants, whether HCQ was present or not (**Fig. 2E**). This observation also indicates that the drug does not impact genome replication. In contrast, infection with infectious Nluc reporter virus reproduced the differential HCQ inhibition as observed in the original non-modified viruses (**Fig. 2F & Fig. 1D**). Taken together, these data indicate that MP4 enters via pH-dependent endocytosis, while MP4-97R/167G entry pathway is independent on endosomal acidification.

**Fig 2.**
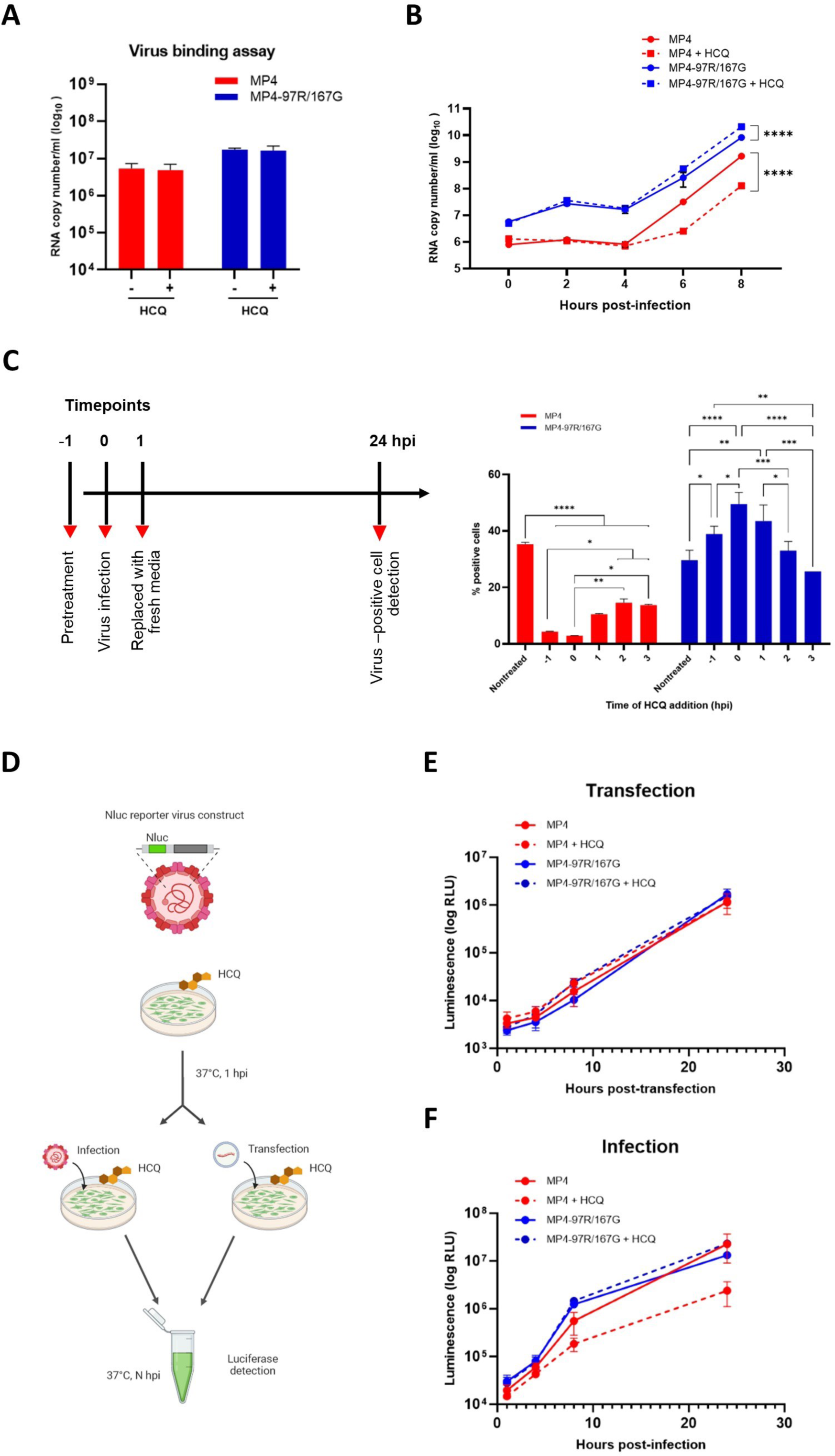
HCQ targets viral entry. (**A**) Virus binding assay in Vero cells in presence of 25 µM HCQ. (**B**) Single-cycle replication kinetic in nontreated and HCQ-treated Vero cells. At each timepoint, cell lysates were collected, and viral RNA copy numbers were quantitated using RT-qPCR (**C**) Time-of-addition assay in Vero cells treated with HCQ starting at different timepoints. Infected cells were quantitated 24 hpi by IF. (**D**) Schematic diagram of Vero cells pre-treated with HCQ and subsequently subjected to transfection of *in vitro* RNA transcript or infection with EV-A71 nanoluciferase (Nluc) reporter viruses. At the indicated timepoints, cell supernatants were collected, and luciferase activity was measured. (**E & F**) Results are expressed in % relative light unit (RLU) of treated versus nontreated virus at indicated timepoints. The mean and S.E.M from biological triplicates are shown. Statistical significance was calculated using two-way ANOVA, comparing treated and untreated control. *p <0.05, **p < 0.01, ***p < 0.001, ****p < 0.0001.

### HCQ differentially impacts MP4 and MP4-97R/167G uncoating

To further dissect the mechanism of action of HCQ on the entry of each variant, we next examined the effect of the drug on viral uncoating. We generated neutral red labelled virus stocks and performed neutral red uncoating assay as previously described^22^. Briefly, viral stocks were produced in presence of neutral red in the dark to induce co-encapsidation of viral genome and neutral red inside viral particles. Photoactivation of neutral red induces the dye to cross-link viral genomes to the capsid and subsequent blockade of viral uncoating^22^. Accordingly, upon infection with neutral red labelled viruses, light exposure only inactivates viruses that have not yet completed uncoating, while virus genomes already released in the cytoplasm are unaffected. This allows to precisely define the timepoint of viral uncoating. Cells were pre-treated with HCQ and then incubated with neutral red-labelled viruses for 1 hr at 37°C for infection. Light inactivation was performed at selected timepoints post-infection, and infected cells were quantified 24 h later by immunostaining (**Fig. 3A**). In the absence of HCQ, the majority of viruses had undergone uncoating between 2 and 3 hpi for both variants (30-70% of uncoated viruses for MP4 and 60-80% for MP4-97R/167G, respectively) (**Fig. 3B**). In the presence of HCQ, MP4 uncoating was completely blocked, even when photoactivation was performed at 4 hpi (**Fig 3B, left panel**). In contrast, the uncoating rate was not inhibited in the presence of HCQ for MP4-97R/167G (**Fig. 3B, right panel**) and the final viral yield was even increased as already observed in **Fig. 1D**.

**Fig 3.**
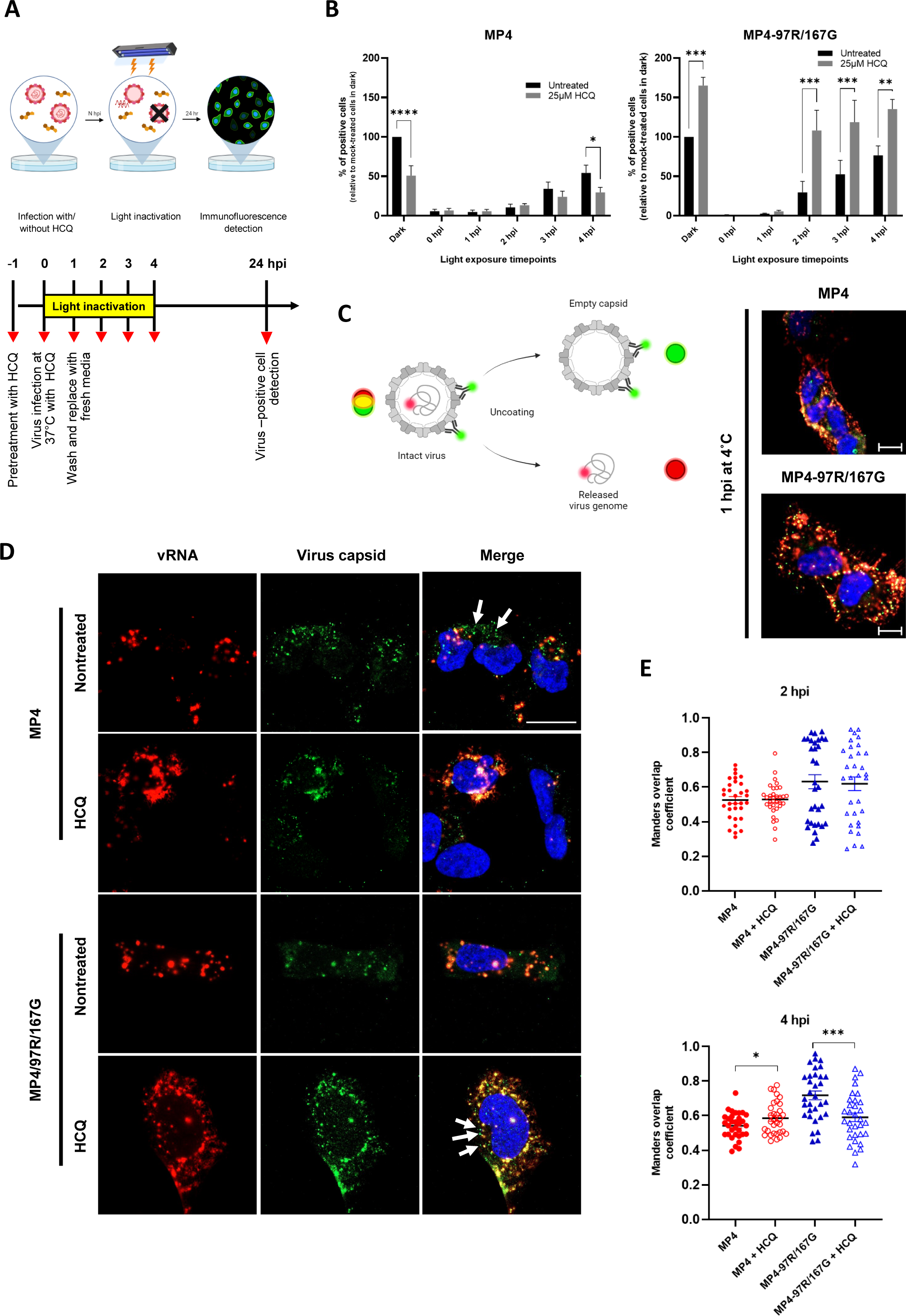
HCQ delays the uncoating of MP4. (**A**) Schematic illustration of the neutral red assay workflow. Vero cells were pre-treated with or without HCQ for 1hr. Neutral red-labelled viruses were allowed for cell infection at 37°C for 1hr. Upon infection, the inoculum was removed and replaced with fresh media. Infected cells were exposed to light for 30 min at different timepoints and further incubated up to 24hpi for IF staining **(B)** Effect of light inactivation on replication of neutral red-labelled MP4 (left panel) or MP4-97R/167G (right panel). Results are plotted as % of virus-positive cells relative to non-treated dark control. Mean and S.E.M of biological triplicates are shown. Statistical significances (two-way ANOVA) were calculated between treated and nontreated conditions. **(C)** Schematic illustration of virus uncoating monitored with the combinational use of RNA-FISH to detect EV-A71 RNA (red) and IF with anti-VP2 Ab to detect the viral capsid (green). Co-staining highlights intact viruses in yellow while empty capsids and free RNA are in green and red, respectively. Representative images (scale, 20 µm) of MP4 and MP4-97R/167G binding after 1hr at 4°C (C, right panel) and of vRNA (red) and capsids (green) with and without HCQ treatment at 4hpi (**D**). Arrows: empty capsid. **(E)** Co-localization of capsid and vRNA in individual cells at 2 hpi and 4 hpi analysed using Mander’s overlap coefficient (n = 32 individual cells from two independent experiments). Statistical comparison (unpaired t-test) of untreated and treated groups. *p < 0.05, **p < 0.01, ***p < 0.001, ****p < 0.0001.

To further validate these results, we then combined fluorescent *in situ* hybridization (FISH) of viral genomic RNA and immunofluorescence staining of viral capsid at early timepoints. Full particles are characterized by colocalization of virus genomic RNA (vRNA) and viral capsid (as shown in 1 hpi at 4°C), while the colocalization is lost following the uncoating process (**Fig. 3C**). Quantification of vRNA and capsid colocalization highlighted no significant difference between the two variants at 2 hpi in presence or absence of HCQ (**Fig. 3E**). However, at 4 hpi (prior to the initiation of replication, see **Fig. 3B**), MP4 uncoating appeared to be inhibited by HCQ, as highlighted by a decrease of empty capsids and an increase of capsid/RNA colocalization in presence of the drug (**Fig. 3D & E**). An opposite effect was observed for MP4-97R/167G, with a reduced capsid/RNA colocalization in the presence of HCQ, indicating that more viruses had undergone uncoating in the presence of HCQ at this time point. Altogether these data indicate that MP4-97R/167G can uncoats at neutral pH and that acidification is instead increasing its replication capacity, while MP4 needs acidic pH to uncoat.

### MP4 relies on late endosomes for uncoating, whereas MP4-97R/167G undergoes uncoating in early endosomes

HCQ is known to inhibit endosomal acidification by accumulating in endosomes in a protonated form. This accumulation leads to endosomal swelling and inhibition of fusion between endosomes and lysosomes within cells, as previously described^23,24^ and as shown in **Fig. 4A**. We thus hypothesized that the two variants could exploit different entry routes, which could explain the different sensitivity to HCQ. We showed that MP4 needs acidic pH to uncoat and is thus expected to release its RNA in late endosomes/lysosomes. In contrast, MP4-97R/167G can uncoat in absence of pH acidification and accordingly in a non-acidic environment. To test this hypothesis, we infected Vero cells transiently expressing a variant of small GTPase Rab5a, a protein involved in the maturation of early endosomes (EE) into late endosomes (LE). This Rab5a-Q79L mutant is constitutively active (CA) and blocks LE maturation (**Fig. 4B**). Viral capsids of MP4 and MP4-97R/167G were observed within EE in both Rab5a WT and CA-expressing cells at 0.5 hpi and 2 hpi (**Fig. S3 and 4C**). However at 7 hpi, the percentage of cells stained for double stranded RNA (dsRNA), a marker of virus replication, was significantly reduced for MP4 in Rab5a CA-expressing cells but not for MP4-97R/167G (**Fig. 4D**). This indicates that MP4-97R/167G genomes were successfully released in the cytoplasm to undergo translation and replication, even in absence of EE fusion to LE. Conversely, a transition from EE to LE with a gradual pH decrease is necessary for MP4 to release its genome.

**Fig 4.**
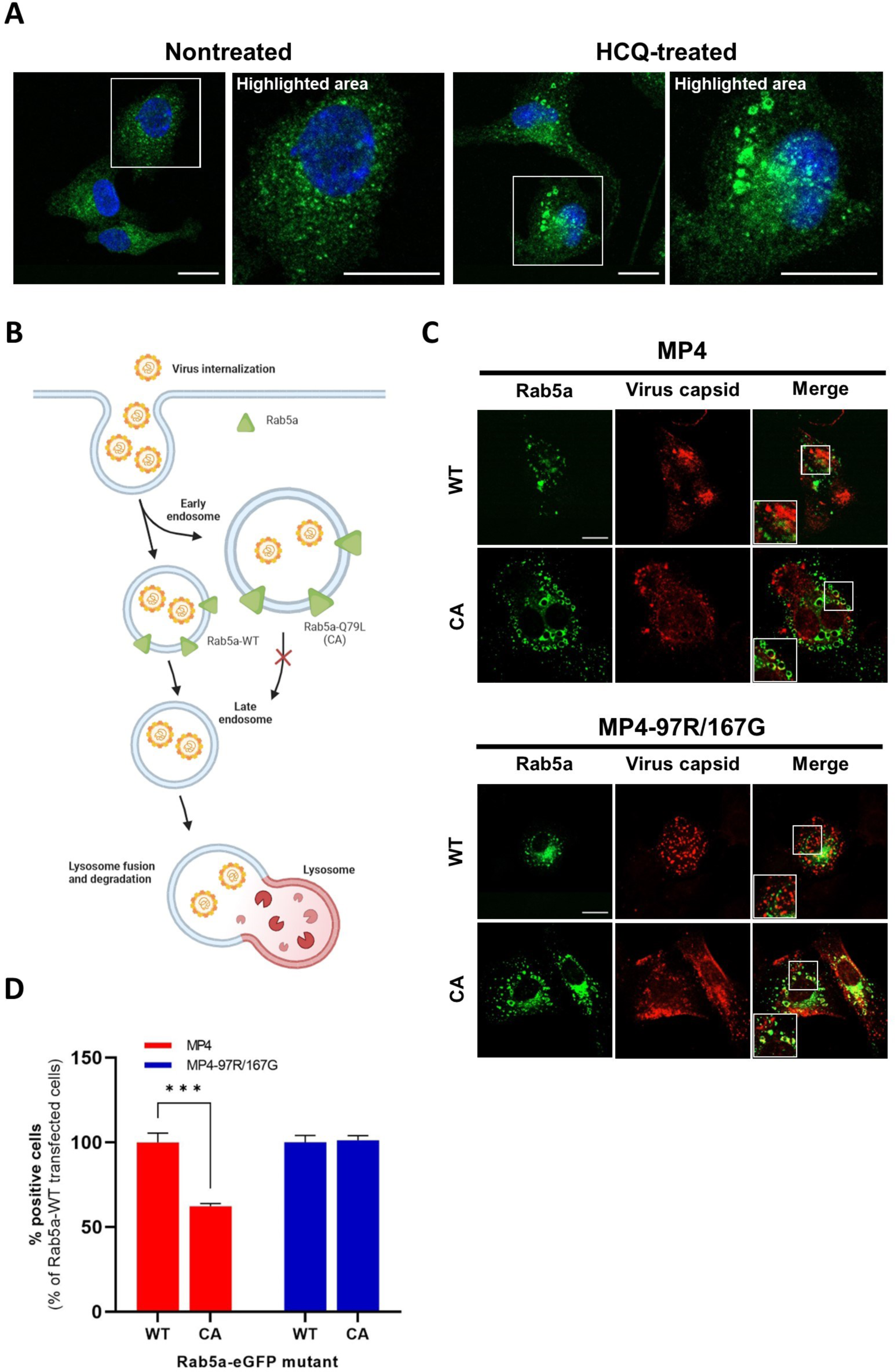
MP4-97R/167G uncoats from early endosomes. (**A**) Nontreated and HCQ-treated Vero cells were stained with anti-EEA-1 antibody (green) to label early endosomes, and DAPI (blue) to label cell nuclei. **(B)** Schematic representation of endosomal route upon overexpression of Rab5a WT or CA mutant. **(C)** Viral capsids (anti-VP2 Ab) localise in early endosomes, 2 hpi of Vero cells transiently expressing Rab5a (in red) WT and CA. **(D)** % of cells stained positive with the anti-dsRNA J2 Ab in FACS-sorted Rab5a-eGFP-expressing cells 7 hpi. Results and statistical significance (two-way ANOVA) are expressed relative to Rab5a WT-expressing cells. Mean and S.E.M from triplicates are shown. ***p < 0.001. In B and D, white boxes are enlarged in the right panel. Scale bar: 20 µm.

### VP1-L97R/E167G substitutions confer affinity for HS but decrease capsid stability

Our data highlighted that both MP4 and MP4-97R/167G enter via a SCARB2-dependent pathway, localize in early endosomes at early times post-infection but exhibit distinct sensitivities to HCQ, a feature independent of their differential use of HS as attachment receptor. We therefore speculated that the varying pH-dependency may be attributed to differences in virion stability. We used DynaMut server^25^ to assess the relative influence of VP1-97R and VP1-167G mutations on interaction networks in their respective local environments (**Fig. S4A**) as well as the impact on overall capsid stability (**Fig. S4B**). Indeed, VP1-L97R mutation is predicted to reduce hydrophobic interactions between the original leucine residue on this position and its neighbors VP1-245Y and VP1-246P. Further, this residue is in close proximity to VP1-244K, thereby adding more positive charge to this site. On the other hand, VP1-E167G mutation causes a loss in hydrogen bonding capacity to VP1-165S and reduces the net negative charge. Altogether, DynaMut analysis predicted reduced interaction capabilities by the substituted residues and changes in electrostatic properties within the region. This is consistent with analyses of vibrational entropy change, indicating that the presence of two mutations results in enhanced local dynamics, which has previously been correlated with reduced capsid stability (**Fig. S4B**)^26,27^.

We therefore speculated that MP4-97R/167G mutant features a lower stability and could bypass the need for acidic pH for uncoating. To test this hypothesis, we subjected these variants to neutral and acidic conditions and assessed their virion structure using negative staining electron microscopy (nsEM) (**Fig. 5A & B**). At acidic pH, there was no notable alteration in the capsid morphology of MP4, which maintained a stable particle diameter of ∼31-33 nm across both pH conditions. On the other hand, for MP4-97R/167G, both 2D images and 3D reconstructions highlighted a loss of density at the center of the viral particles, as well as an expansion in size for a subset of particles (diameter ranging from 31 to 41 nm at pH5 versus 31 to 33 nm at pH7), indicating partial virus uncoating. We then performed temperature sensitivity assay by heating viruses at different temperatures for 1 h before inoculation on cells. Quantification of infected cells at 24 hpi further confirmed that MP4 capsid is more resistant to higher temperature as 80% of MP4 population survived a 50°C thermal stress compared to only 50% for MP4-97R/167G (**Fig. 5C**). Consistently, the predictions of Gibbs free energy change (ΔΔG) induced by these mutations further supported that both mutations induce slight destabilization of the capsid structure, regardless of pH and temperature (**Table S1**).

**Fig 5.**
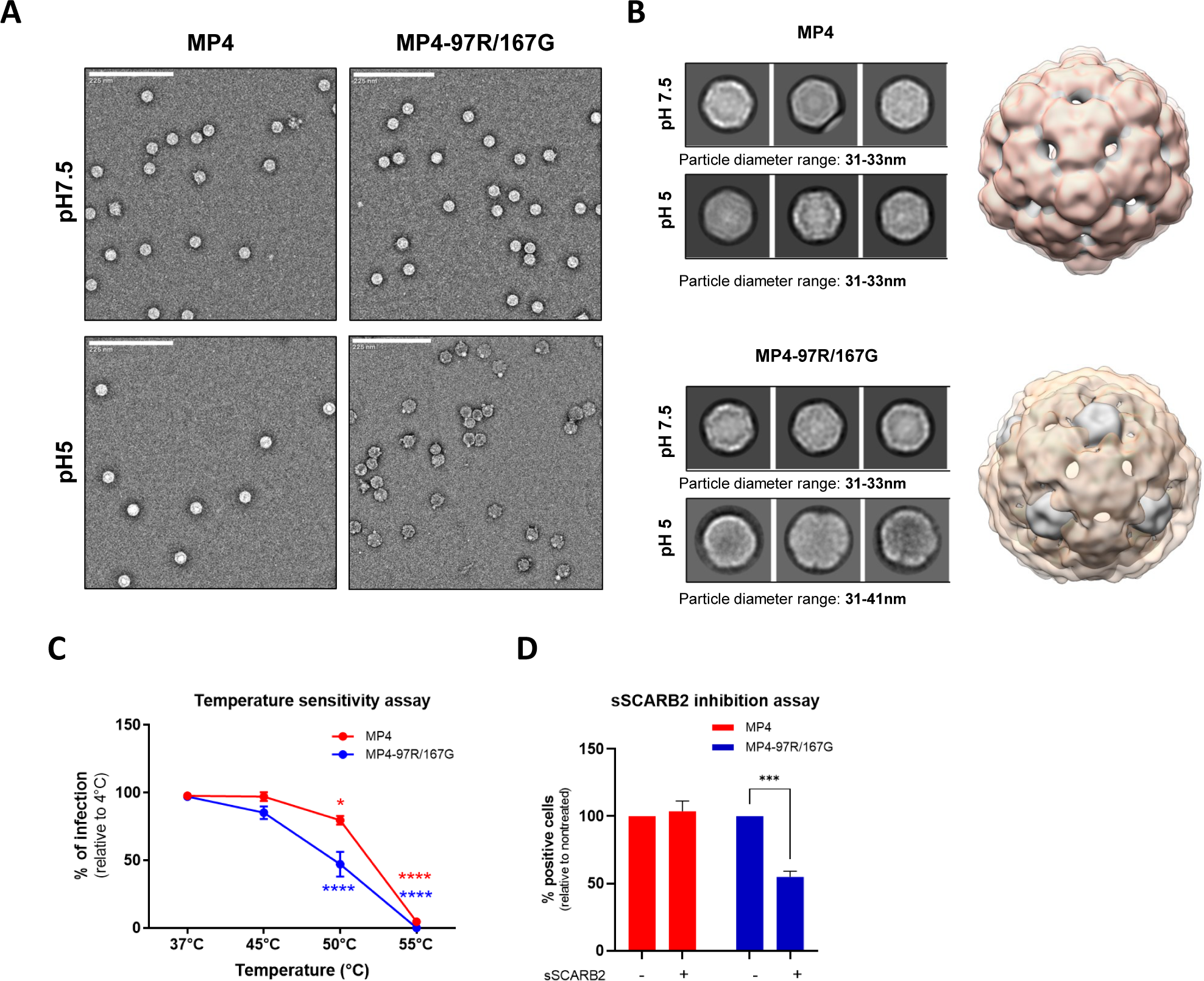
MP4 displayed stronger capsid stability and reduced sensitivity to acidification and high temperatures. (**A**) nsEM analysis of MP4 and MP4-97R/167G incubated at pH7 and pH5. Representative raw micrographs are shown in each case. (**B**) Representative 2D class averages generated from datasets shown in panel A (box size = 54nm; left) and the overlay of the corresponding 3D maps (right). Grey and orange shade indicates virus particle reconstructions at pH7 and pH5, respectively. (**C**) Temperature sensitivity assay. Infected Vero cells were quantitated by immunostaining with an anti-VP2 Ab at 24 hpi after 1 hr incubation at increasing temperatures. Results are shown as % of virus-positive cells relative to 4°C treated control. Error bars indicate mean and S.E.M from biological triplicates. (**D**) For sSCARB2 inhibition assay, viruses were incubated 1h at 37°C with 1µg of soluble SCARB2 (sSCARB2) before infection of Vero cells. Infected Vero cells were quantitated by immunostaining with an anti-VP2 Ab at 24 hpi. Results are shown as % of virus-positive cells relative to nontreated controls. Statistically significance was calculated with two-way ANOVA. ***p < 0.001, ****p < 0.0001.

As MP4-97R/167G is less stable, we hypothesized that binding to SCARB2 may be sufficient to trigger its capsid opening. We conducted a competitive experiment and compared the infectivity of the two variants after incubation with soluble SCARB2 (sSCARB2) at neutral pH for 1 hr at 37°C. We observed that MP4-97R/167G but not MP4 variant lost infectivity upon pre-exposure to sSCARB2 (**Fig. 5D**). Altogether, our results confirmed that MP4-97R/167G capsid is less stable and highly sensitive to thermal and acidic stresses as well as receptor binding, which are sufficient triggers to initiate virus capsid disruption and subsequent viral uncoating.

### Resistance to HCQ and reduced capsid stability extend to other strong heparan sulfate-binding strains

To check if our observations could extend to other HS-binding variants, we checked the effect of mutation present at position VP1-145, a residue known to play a key role in modulating viral HS-binding capacity and *in vivo* virulence. EV-A71 variants (sub-genogroup B4) with wild-type VP1-145E is a weak HS binder and is attenuated in mice while the cell-adapted VP1-145Q variant is a strong HS-binder and virulent in mice^11^. As shown in **Fig. 6A**, HCQ enhanced infectivity of VP1-145Q variant but reduced infectivity of VP1-145E variant, consistent with what we observed with the MP4-97R/167G. Both temperature sensitivity **(Fig. 6B)** and sSCARB2 inhibition assays **(Fig 6C)** also confirmed that the capsid of VP1-145E variant is much more stable compared to VP1-145Q variant, in line with free energy change prediction as shown in **Table S1**. These data further reinforced the observation that HS-binding phenotype is inversely correlated with virus capsid stability.

**Fig 6.**
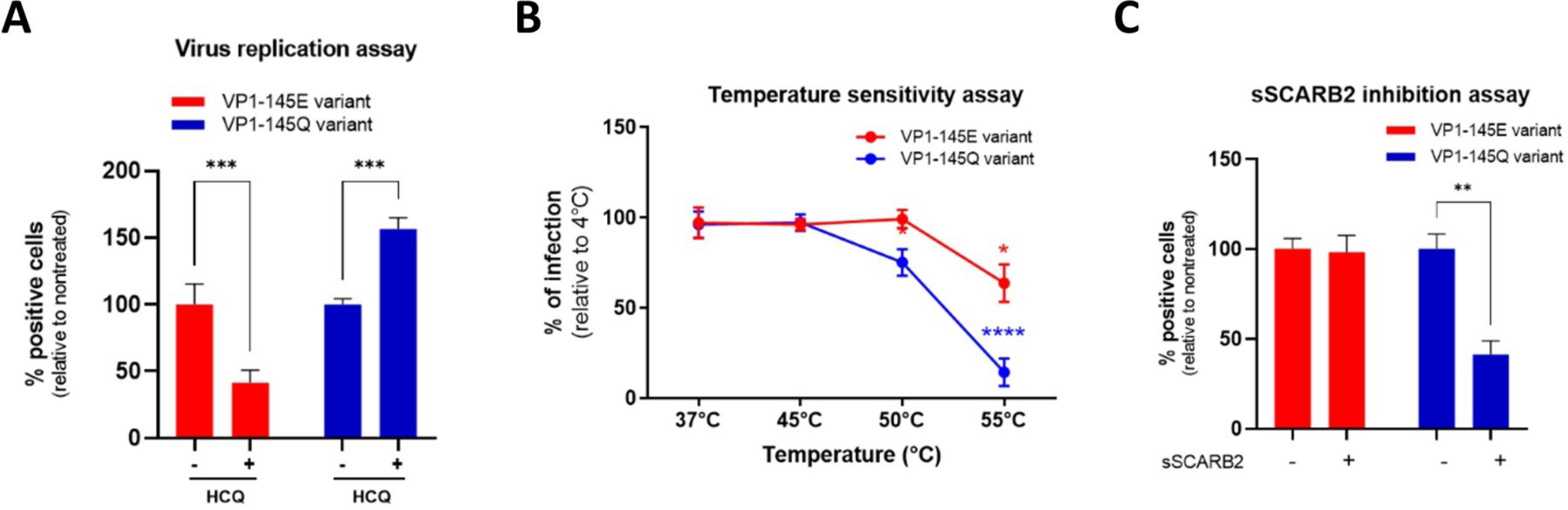
Heparan-sulfate-binding VP1-145Q variant exhibits resistance to HCQ and higher sensitivity to sSCARB2 inhibition and thermal stress. (**A**) Virus inhibitory assay with VP1-145 variants were performed with 25 µg HCQ on Vero cells. **(B)** For temperature sensitivity assays, VP1-145 variants were incubated at increasing temperature for 1hr before inoculated onto Vero cells. **(C)** For sSCARB2 inhibition assay, VP1-145 variants were incubated 1 h at 37°C with 1 µg of soluble SCARB2 (sSCARB2) before infection of Vero cells. Infected cells were quantitated by immunostaining with anti-VP2 Ab at 24 hpi. Results are shown as % of virus-positive cells relative to nontreated control (A & C) or 4°C treated control (B). Mean and S.E.M of biological triplicates are shown. Statistically significant differences (two-way ANOVA) are shown. **p < 0.01, ***p < 0.001, ****p < 0.0001.

## Discussion

Acidic pH is an important trigger for viral uncoating and many enveloped or non-enveloped viruses, including influenza A virus^28,29^, human adenovirus^30^, foot-and-mouth disease virus^31^, Semliki forest virus^32^, that enter the cell through pH-dependent endocytosis^33^. Similarly, for EV-A71, binding to SCARB2 and subsequent endosomal acidification are required for uncoating^4,7^. In this study, we provide new insights on the impact of mutations in the VP1 capsid protein leading to strong HS affinity, on the uncoating process of EV-A71. We show that, unlike the mouse-adapted EV-A71 MP4 strain, the MP4-97R/167G-derived double mutant, which has a high affinity for HS, does not require acidification for uncoating and can release its genome under neutral or weakly acidic environment of early endosomes.

To demonstrate the importance of acidification on both MP4 and MP4-97R/167G variant uncoating, we used two lysosomotropic drugs, namely BAF-A1 and HCQ, that increase endosomal pH by distinct means. On one hand, BAF-A1 inhibits the vacuolar H+ ATPase (V-ATPase), preventing the acidification process and thereby elevating the endosomal pH ^34-38^. On the other hand, HCQ, a less toxic derivative of the antimalarial drug chloroquine, acts as a weak base that can be protonated and trapped in the acidic environment of cellular organelles^39-41^. In addition to this effect, HCQ can impact other cellular pathways, such as autophagy, a cellular process that has been demonstrated to be induced by EV-A71 to create a favorable environment for its replication^42,43^. However, we show here that the differential effects on MP4 and MP4-97R/167G occur during the uncoating process rather than in later stages of the cycle, such as virus genome replication^40^. Our results thus underline that, despite their distinct modes of action, both HCQ and BAF-A1 influenced virus entry through their effect on endocytic compartments, as both compounds ultimately inhibit the reduction in endosomal pH levels. In addition, the fact that differential sensitivity to HCQ was retained even in cells devoid of HS at their surface by treatment with heparinase or sodium chlorate pointed out that this pH-independent mode of entry of MP4-97R/167G is not linked to the use of HS as an attachment receptor. Interestingly, our experiments using the Rab5a CA to block the transition of EE to acidic LE^44,45^, indicate that binding to SCARB2 is sufficient to trigger MP4-97R/167G genome release into the cytosol even in the near neutral pH of the EE (pH ∼6.0 to 6.5), whereas MP4 requires the acidity of LE and/or lysosomes (pH ∼5.0 to 5.5) to uncoat efficiently^46^. These observations led us to hypothesize that the two variants exhibit intrinsic differences in capsid stability. We conducted various tests to compare how each variant reacted to heat, sSCARB2 and low pH, and found that MP4-97R/167G was more sensitive to all these conditions. Particularly, we used nsEM to study the properties of viral particles and observed an expansion of MP4-97R/167G capsid following the exposure to pH 5. These data, plus the virus structural dynamics prediction and free energy change computation, all indicate that MP4-97R/167G presents reduced capsid stability compared to MP4.

One strength of our study lies in the fact that we were able to extend these results to another HS-dependent EV-A71 strain, VP1-145Q variant. Given that these two HS-dependent variants share the characteristic of having incorporated a less acidic amino acid within the VP1 capsid protein, we hypothesized that an increase in positive charges within the capsid not only enhances affinity for HS but also alters capsid stability, consequently impacting the virus entry mechanism. In the same line, a thermostable EV-A71 variant (VP1-K162E, change of a basic to an acidic residue) isolated from serial passages at higher temperatures was shown to be less efficient at uncoating with poorer cell infectivity but more virulent in mice^47^. Interestingly, this variant showed a more expanded conformation compared to the original non-mutated virus^47^. While the thermostable variant showed no difference in binding to SCARB2 receptor, the binding affinity to heparin was greatly reduced, an observation consistent with what we noticed for MP4. Additional experiments with cell-adapted, HS-binding viruses will help to define whether HS-binding is always associated with a loss of virion stability and whether these findings could event extend to other group of viruses. Interestingly, mutations conferring similar *in vitro* phenotypes were observed for other enteroviruses such as rhinovirus A16 (RV-A16) and coxsackievirus B3 (CV-B3). For RV-A16, capsid mutations conferring resistance to endosomal acidification inhibitors also abrogated the need for acidic pH for uncoating. More importantly, these mutations were also associated with higher sensitivity to low pH, high temperatures, and binding to soluble receptors^48^. For CV-B3, a fast-growing variant was shown to exhibit faster genome release and destabilized capsid, and this led to attenuated virulence in mice^49^. This existing literature aligns with our findings, suggesting that alterations in capsid stability due to amino acid changes may significantly impact various aspects of the virion and its life cycle, including sensitivity to environmental factors, receptor interactions, and infection rates.

In the light of these published studies and our data, we propose the following model depicting the relationship between HS-binding, capsid stability and viral fitness *in vitro* and *in vivo* (schematized in **Fig. 7**). Viruses undergo mutations and positive selection to adapt to different environments^50,51^. EV-A71 can take advantage of the high plasticity of its capsid to optimize its fitness upon environmental changes. Many strains adapt to use HS *in vitro* due to the abundant expression of this attachment receptor in cell lines. To do so, they usually acquire additional positively charged amino acids within outward-facing VP1 domains proximal to the capsid 5-fold axis. In addition to help the virus to attach on the cell surface and find the SCARB2, we show here that these mutations concomitantly decrease virion stability. This further contributes to higher multiplication in cell lines by triggering uncoating rapidly after internalisation, within EE, without the need of acidic pH. Interestingly in our experiments, acidification inhibitors improved rather than inhibited viral fitness of MP4-97R/167G. Although additional experiments are required to define the mechanism behind this observation, it could occur via the protection of virions that have not yet uncoated when endosomes fuse with lysosomes. In this context, absence of acidification would improve the chance of these virions to release their genome in the cytoplasm, while exposure to acidic pH would induce viral opening within the late endosomes. The situation may differ significantly *in vivo* as strong HS binders are attenuated. Koike and colleagues have demonstrated that strong binding to HS induces virus trapping *in vivo*^9-11,19^. Our data suggest that the associated decreased stability may further contribute to viral attenuation. To be virulent *in vivo*, a non-enveloped virus must have a sufficiently stable capsid to resist unfavourable environmental conditions, both during dissemination within a host and during transmission between hosts. This last point is particularly important for EV-A71, which is transmitted via the fecal-oral route and must therefore resist the acidic pH of the gastrointestinal tract before reaching the intestinal mucosa, its main multiplication site. Capsid stability thus ensures that virus genome release occurs only in a proper environment, but in turn renders the virus dependent on both SCARB2 and acidic pH for uncoating^4,7^.

**Fig 7.**
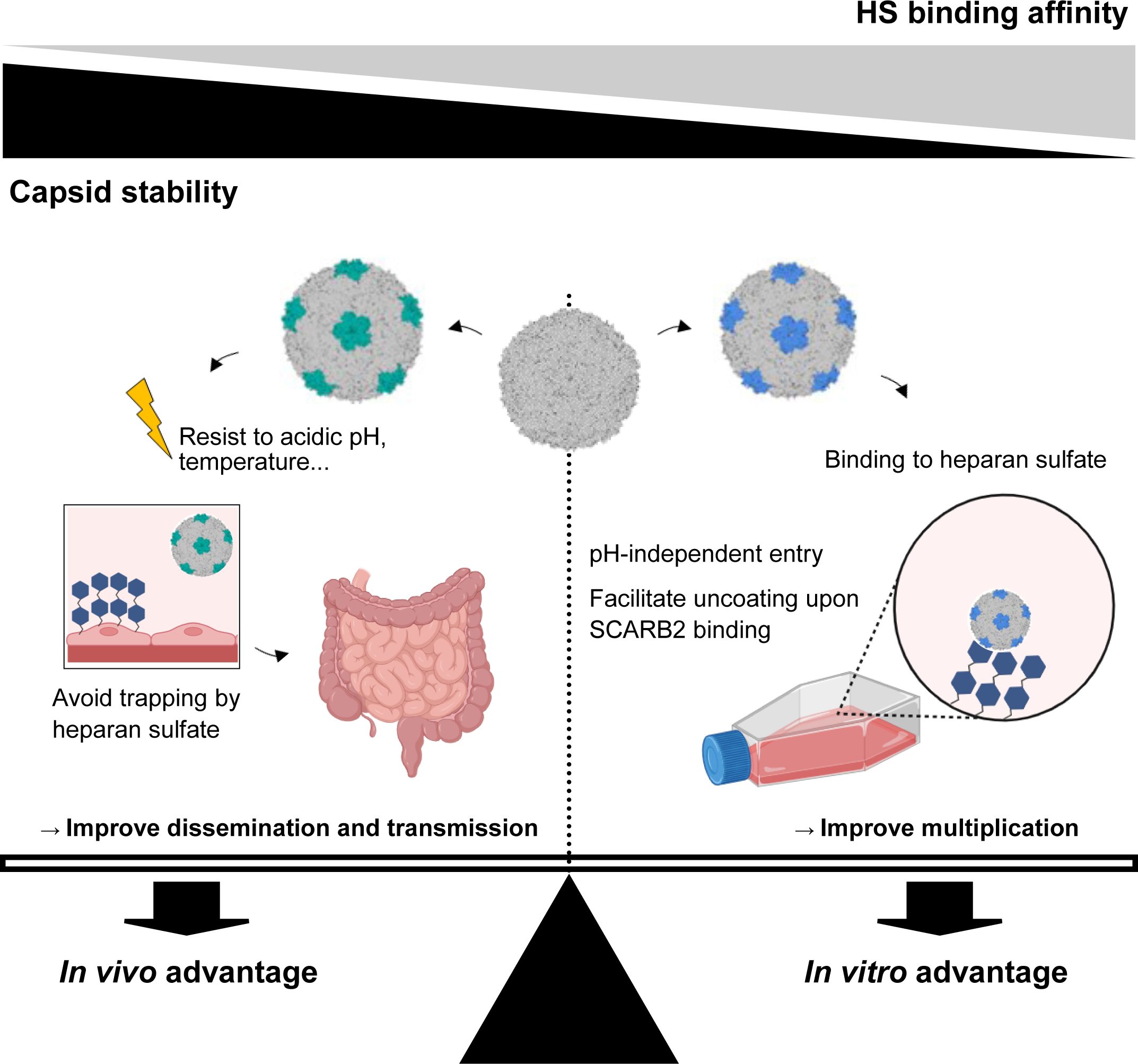
Seesaw model depicting the interplay between capsid mutations, heparan sulfate-binding, capsid stability as well as the resulting fitness changes in both *in vitro* and *in vivo* settings. Viruses undergo continuous mutations to optimize fitness across diverse environments. In cell culture, they adapt to attain an ‘*in vitro* advantage’ by decreasing capsid stability while acquiring HS-binding capacity, consequently enhancing their infectivity. Conversely, during human infection, viruses adapt to secure an ‘*in vivo* advantage’ by bolstering capsid stability, relinquishing HS-binding capacity, and thereby evading viral trapping and resisting environmental stresses.

Of note, although strong HS-binder are clearly attenuated in mice, the situation in humans is more puzzling as strains with Q or G at position VP1-145 have been associated with severe neurological cases and the methodology used in those studies excluded emergence of mutation during cell culture^52,53^. Furthermore, we previously showed that intermediate HS-binding affinity can result in increased virulence even in mouse models^19^. It is still unclear if there is a lack of trapping and/or limited impact on viral stability under these conditions. To conclude, to reach optimal fitness *in vitro* and *in vivo*, the virus needs to find the correct balance between HS binding and capsid stability (**Fig. 7**). Our study improves the current knowledge on the mechanism behind the *in vivo* attenuation of cell culture-adapted viruses. It also opens the doors to new antiviral strategies targeting endosomal acidification as well new principles for vaccine design based on attenuated -acid-independent variants to help combat EV-A71.

## Materials and methods

### Chemical reagents

Chemical reagents used in this study were listed as following: hydroxychloroquine (Tocris), bafilomycin-A1 (InvivoGen), sodium chlorate (NaClO_3_, Sigma-Aldrich), neutral red (Sigma Aldrich) and LysoTracker Deep Red (Thermo Fisher Scientific).

### Cell lines and virus

Vero (monkey kidney; ATCC CCL-81) and human rhabdomyosarcoma cells (RD; ATCC no.: CCL-136) were propagated in Dulbecco’s Modified Eagle Medium (DMEM) and GlutaMAX (31966021, Thermo Fisher Scientific) containing 10% fetal bovine serum (FBS). RD-SCARB2-KO^20^ and RD-ΔEXT1+hSCARB2^54^ cells were maintained in DMEM supplemented with 10 µg/ml puromycin (58-58-2, InvivoGen). All infected cells were maintained in media supplemented with 2.5% FBS. All cells were maintained at 37°C in 5% CO_2_. Viruses used in this study including MP4, MP4-97R/167G (Genbank accession number: JN544419; subgenogroup C2), IEQ (EV-A71 VP1-145Q variant; Genbank accession number: AF316321; subgenogroup B4) and IEE (EV-A71 VP1-145E variant) strains were prepared as previously described^11,19^. For Nluc reporter virus, Nluc gene was inserted between 5’ UTR and VP4 of the virus as previously described^55^. Viruses were generated in RD-ΔEXT1+hSCARB2 cells^54^, propagated for an additional passage and used as working stocks. All virus stocks were sequenced for confirmation (Fasteris) prior to experiments.

### Plasmids

Plasmids encoding eGFP-Rab5a and eGFP-Rab5a Q79L are kind gifts from Pierre-Yves Lozach (University Claude Bernard Lyon 1). Both IEQ and IEE plasmids (Genbank accession number: JN544419: AF316321; subgenogroup B4) strains are kind gifts from Yoke Fun Chan (University of Malaya).

### Virus inhibitory assay and time-of-addition assay

For virus inhibitory assay, cells were pre-treated either with drugs for 1 hour at 37 °C. Viruses (MOI 0.1) were inoculated onto cells in presence of drugs for 1 hour at 37 °C. Upon infection, inocula were removed and cells were rinsed thoroughly with phosphate buffered saline buffer (PBS) before incubated with fresh media up to 24 hpi. For time-of-addition assay, HCQ was either pretreated (-1hpi), introduced during virus infection (0hpi) or introduced onto cells at post-infection (1, 2 and 3hpi) for 1hr. After incubation, cells were rinsed with PBS before loaded with maintenance media and incubate up to 24 hpi. For both assays, infected cells were fixed for immunofluorescence staining for virus-positive cells detection.

### Virus binding and replication assay

All the experiments were done on Vero cells seeded in 96 wells plate. For virus binding assay, cells were incubated with 1 × 10^8^ RNA copy number/ml virus for 1 hour at 4 °C. The inocula were removed and rinsed with cold PBS twice, and then subjected to cell lysis for RNA extraction and qRT-PCR quantitation. For virus replication assay, cell monolayers were incubated with virus for 1 hour at 37 °C. The inocula were removed, rinsed with PBS, and then further incubated up to 24 hpi at 37 °C. Infected cells were lysed and viral RNA was quantified by qRT-PCR.

### Heparan sulfate removal

Both enzymatic and chemical methods, heparinase assay and sodium chlorate (NaClO_3_) treatment respectively, were used to cleave HS from cell surface. For heparinase assay, the cells were first rinsed with PBS and then incubated with 3.5 mIU/ml of heparinase III (AmsBio) diluted in 0.1 M sodium acetate pH 7.0, 1 mM calcium acetate and 0.2% BSA for 1 hour at 37 °C. Meanwhile, mock-treated cells were incubated with heparinase buffer. Upon incubation, cells were washed twice prior to virus infection. For NaClO_3_ treatment, cells were propagated in presence of 30 mM NaClO_3_ at least one passage before experiment. The cells were also pre-seeded in media supplemented with NaClO_3_ and incubated overnight before the experiment.

### Immunofluorescence and confocal imaging

To detect infected cells, cells were fixed with absolute methanol (Sigma Aldrich) at room temperature for 10 min and then incubated with blocking buffer consisting of 5% BSA (PanReac Applichem) and 0.05% TritonX-100 (PanReac Applichem) for 20 min. Fixed cells were first incubated with anti-EV-A71 capsid monoclonal antibody MAB979 (1:1000; Sigma) for 1 hour at 37 °C and then with Alexa Fluor 488-conjugated secondary antibodies (1:2000; Thermo Fisher Scientific) dissolved in DAPI solution for 1 hour at 37°C. To detect dsRNA, infected cells were fixed with 4% paraformaldehyde (Santa Cruz) and incubated with blocking buffer, cells were incubated with anti-dsRNA monoclonal antibody J2 (1:500; Scicons) for 1 hour at 37 °C and then with Alexa Fluor 594-conjugated secondary antibodies (1:2000; Thermo Fisher Scientific) dissolved in DAPI solution for 1 hour at 37°C. Stained cells were acquired using ImageXpress Pico (Molecular Devices) and percentages of positive cells were determined using CellReporterXpress software. For confocal imaging, immunofluorescence staining was performed with EE and lysosomes were stained using anti-EEA1 (1:100; Santa Cruz) and anti-LAMP1 (1:100; Cell Signalling), respectively, for 1 hour at 37 °C and then with Alexa Fluor 488-conjugated secondary antibodies (1:200) dissolved in DAPI solution for 1 hour at 37°C. The stained slides were mounted under coverslip (Hecht Assistent) with Fluoromount G mounting medium (Southern Biotech) and analyzed using Zeiss LSM 800 confocal microscopy.

### RNAscope FISH detection and colocalization experiments

For FISH, cells were seeded on Nunc LabTek II chamber slides (Thermo Fisher Scientific) and fixed with 4% paraformaldehyde. To detect viral RNA in infected cells, fixed cells were processed for RNAscope FISH using RNAscope Multiplex Fluorescent V2 assay (Biotechne) according to manufacturer’s protocol. In brief, the cells were hybridized with V-EV71-C1 probe (Biotechne) at 40°C for 2 hours and then the signals were revealed using TSA Vivid 570 kit (Tocris). The slides were then incubated with blocking buffer and incubated with MAB979 (1:100) at 4°C overnight, followed by incubation with Alexa Fluor 488-conjugated secondary antibodies (1:200) at room temperature for 30min. After incubated with DAPI for 30s, the stained slides were mounted under coverslip with mounting medium and analyzed using Zeiss LSM 800 confocal microscopy. Images were acquired and analysed using ZEN 3.2 software.

### Luciferase assay

Enterovirus-A71 nanoluciferase (Nluc) reporter particles were used to study virus replication bypassing cell entry in presence and absence of drug. Briefly, the reporter virus plasmid was linearized and *in vitro* transcribed to generate RNA using T7 RiboMax Express Large Scale RNA Production System (Promega). Transcribed RNA was purified using RNeasy Mini Kit (Qiagen) and then transfected in RD cells using Lipofectamine 2000 (Thermo Fisher Scientific). At certain timepoints, cell supernatants were harvested for luciferase activity detection using Nano-Glo Luciferase Assay System kit (Promega) on Glomax Multi-Detection System (Promega).

### Neutral red uncoating assay

To generate neutral red (NR)-labelled viruses, virus stocks were propagated in cells in presence of 5 µg/ml neutral red (Aldrich). The virus stocks were harvested at 3 dpi and titered. For uncoating assay, NR-labelled viruses were infected at 37 °C for 1 hr in the dark then washed twice with PBS and loaded with FluoroBrite DMEM (Thermo Fisher Scientific) supplemented with 2.5% FBS. At certain timepoints, infected cells were exposed to light for 30 min and then allowed to incubate up to 24 hpi. Infected cells were analysed using immunofluorescence as stated earlier.

### Virus infection in Rab5a-transfected cells

Vero cells (1.5 ×10^6^) were transfected with 25µg of Rab5a-eGFP plasmids using Lipofectamine 3000 (Thermo Fisher Scientific). The next day, transfected cells were harvested, resuspended in buffer (PBS, 2nM EDTA, 1% BSA), and subjected to fluorescence-activated flow cytometry (FACS) on S3 Cell Sorter (Biorad). EGFP-positive cells were sorted, collected and then further propagated at least one day before virus infection. For virus infection, cells were infected with virus (MOI 1.5) for 1 hour at 37°C. The inocula were removed, rinsed with PBS, and cells were further incubated up to 7hpi at 37 °C. Cells were then stained with anti-dsRNA as described above.

### Temperature sensitivity assay and shSCARB2 inhibition assay

Viruses (MOI 0.5) were incubated at different temperatures (4°C, 37°C, 45°C, 50°C and 55°C) for 1hr. Upon incubation, viruses were immediately transferred onto ice for cooling down before inoculated onto cells for 1hr at 37°C. Cells were washed and allowed to incubate in maintenance media up to 24hpi before virus-positive cell detection using immunofluorescence. For SCARB2 inhibition assay, viruses were incubated with 1 µg of soluble recombinant human SCARB2-FC chimera protein (bio-techne) at 37°C for 1hr. The mixture was then inoculated onto cells at 37°C for 1hr. Upon incubation, cells were washed, and allowed to incubate in maintenance media up to 7hpi before lysed the cells for viral RNA quantitation.

### Electron microscopy (EM)

For structural analyses, virus stocks were first inactivated by formaldehyde treatment. Formaldehyde at 100 µg/ml final concentration was added to the virus stock and incubated at 37°C for 3 days. Inactivated viruses were purified through 30% sucrose cushion at 32,000 rpm in SW32 Ti rotor (Beckman Coulter) for 14 hr at 4°C, followed by sedimentation through a discontinuous 20-45% (w/v) sucrose at SW41 Ti rotor (Beckman Coulter) for 12 hr at 4°C. The purified stocks were then subjected to HiPrep 16/60 Sephacryl S-500 HR column (Sigma Aldrich) with 25 mM Tris-HCl + 150 mM NaCl (pH 7.5) as the running buffer. Fractions corresponding to EV A71 particles were pooled and concentrated to 0.3 – 1.1 mg/mL using Amicon Ultra centrifugal filter units with 100 kDa cutoff (Millipore Sigma). For pH-based assays we prepared Tris-Acetate-based buffers at pH 5 and pH 7.5. The buffers comprised 150 mM NaCl and a 100 mM mix of Tris base and acetic acid at the ratio necessary to reach the desired pH. Each EV A71 variant was diluted to 100 µg/ml in the two buffer and incubated for 30 minutes. Following incubation, the samples were applied onto negative stain EM grids (Cat # CF300-Cu-50, Electron Microscopy Sciences). Prior to sample application the grids were glow discharged for 30 seconds. 2% solution of uranyl formate was used for staining. The grids were imaged on a Talos L120C G2 microscope (Thermo Fisher Scientific) running at 120 kV and featuring the CETA 4k camera. EPU software from Thermo Fisher Scientific was used for data acquisition, and all data processing was performed in the cryoSPARC package^56^. Each dataset comprised 100-200 micrographs, and 2’000-10’000 virus-corresponding particles. Particles were extracted from micrographs and subjected to 2D classification. 3D reconstruction was performed using Ab initio algorithm with icosahedral symmetry imposed.

### Computational analysis of virus capsid protein structure stability

To assess the virus capsid protein structure stability, EV-A71 crystal structures with PDB ID of 3J22 and 4AED were used for MP4 and VP1-145 variants, respectively. I-mutant 2.0 server^57^ was used to predict the free energy stability change upon introduction of mutation into virus capsid VP1 protein. Visualization of mutational effects on interatomic interactions and prediction of molecule flexibility were performed on DynaMut server^25^.

### Schematic diagram and statistical analysis

All schematic diagrams and illustrations were created via BioRender.com. All data and statistical analyses were generated using GraphPad Prism 9. All drug treatment experiments were analyzed with one-way and two-way ANOVA. For dose-dependent inhibitory assay, area under curve (AUC) was calculated and analyzed using one-way ANOVA. Degree of colocalization of virus capsid and vRNA in individual cells was measured using Mander’s overlap coefficient calculation in ZEN 3.2 software. Data were presented as mean ± SEM. *p < 0.05, **p < 0.01, ***p < 0.001, ****p < 0.0001. and not significant (n.s.).

## Acknowledgement

This work was funded in part by the Swiss national foundation (Grant N° 310030_184777 to CT) and by the University of Geneva (Salary to HKT). We would like to thank Prof Satoshi Koike and Dr Kyousuke Kobayashi from Tokyo Metropolitan Institute of Medical Science, Japan for providing RD-ΔEXT1+hSCARB2 cells, Prof Jen-Ren Wang from National Cheng Kung University, Taiwan for providing infectious clone plasmids EV-A71/MP4, Prof Pierre-Yves Lozach from Université Claude Bernard Lyon 1 for providing plasmids encoding eGFP-Rab5a and eGFP-Rab5a Q79L, Prof Yoke Fun Chan from University of Malaya for providing IEQ and IEE infectious clone plasmids. We would also like to acknowledge Jessica Swanson, Dr Natalie Kingston and Prof Nicola Stonehouse for giving advice and guidance about virus purification. Electron microscopy data was collected at the Interdisciplinary Centre for Electron Microscopy (CIME) at EPFL with assistance from Davide Demurtas, PhD. Electron microscopy data was processed using the computational infrastructure provided by the IT department of the School of Life Sciences (SV-IT) at EPFL. The authors express sincere gratitude to the CIME and SV-IT personnel for their contribution.

## Figure legends

**Fig S1.**
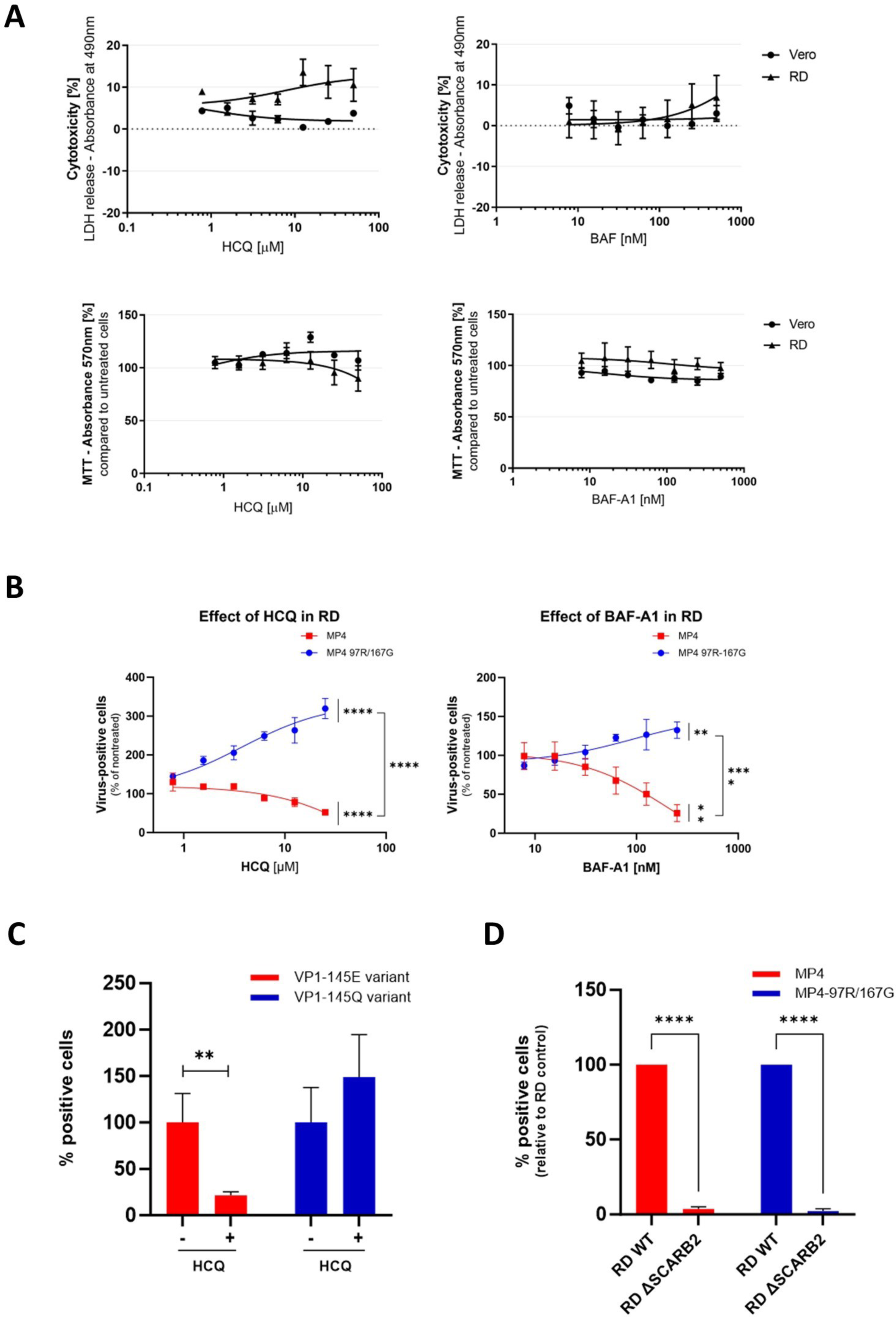
Lysosomotropic drugs nontoxic dose-range and differential inhibition of HS-dependent and independent variants. (**A**) Cytotoxicity effect of lysosomotropic drugs evaluated with LDH and MTT assays. RD and Vero cells were treated with a range of different concentrations of HCQ or BAF-A1 for 2 hr. At 24 hours post-treatment, cell supernatants and lysates were collected for LDH assay and MTT assay, respectively, to determine cytotoxicity effect (n =2). (**B**) Dose response assay with HCQ and BAF-A1 on RD cells were performed exactly like in Vero cells (**Fig.1**). Infected cells (stained with anti-VP2 Ab) were quantitated at 24 hpi after treatment with increasing drug concentrations. Results are shown as % of virus-positive cells relative to nontreated control. AUC was calculated and statistical significance (one-way ANOVA) between treated and untreated virus or between treated MP4 and MP4-97R/167G are shown. Mean and S.E.M of biological triplicates are shown. *p < 0.05, **p < 0.01, ***p < 0.001, ****p < 0.0001.

**Fig S2.**
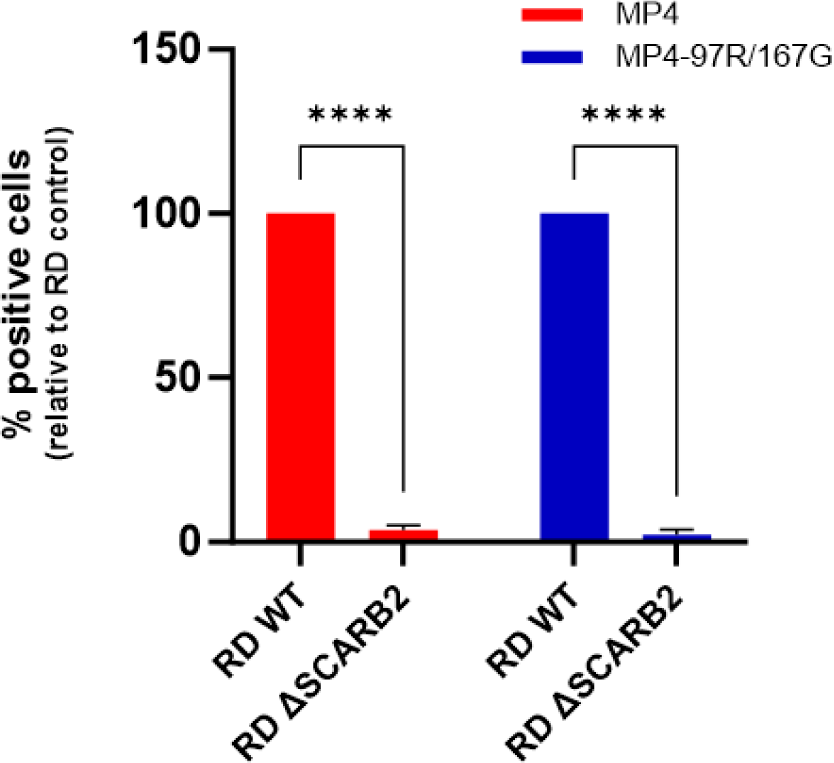
Both EV-A71 variants are strictly dependent on SCARB2 for infection. Virus infection was performed on RD WT and RD ΔSCARB2 cells. Cells were lysed, and viral RNA copy numbers were quantitated at 24 hpi using RT-qPCR. Results are expressed as % Virus RNA copy number relative to RD WT cells (set to 100%). Mean and S.E.M of biological triplicates are shown. ****p < 0.0001.

**Fig S3.**
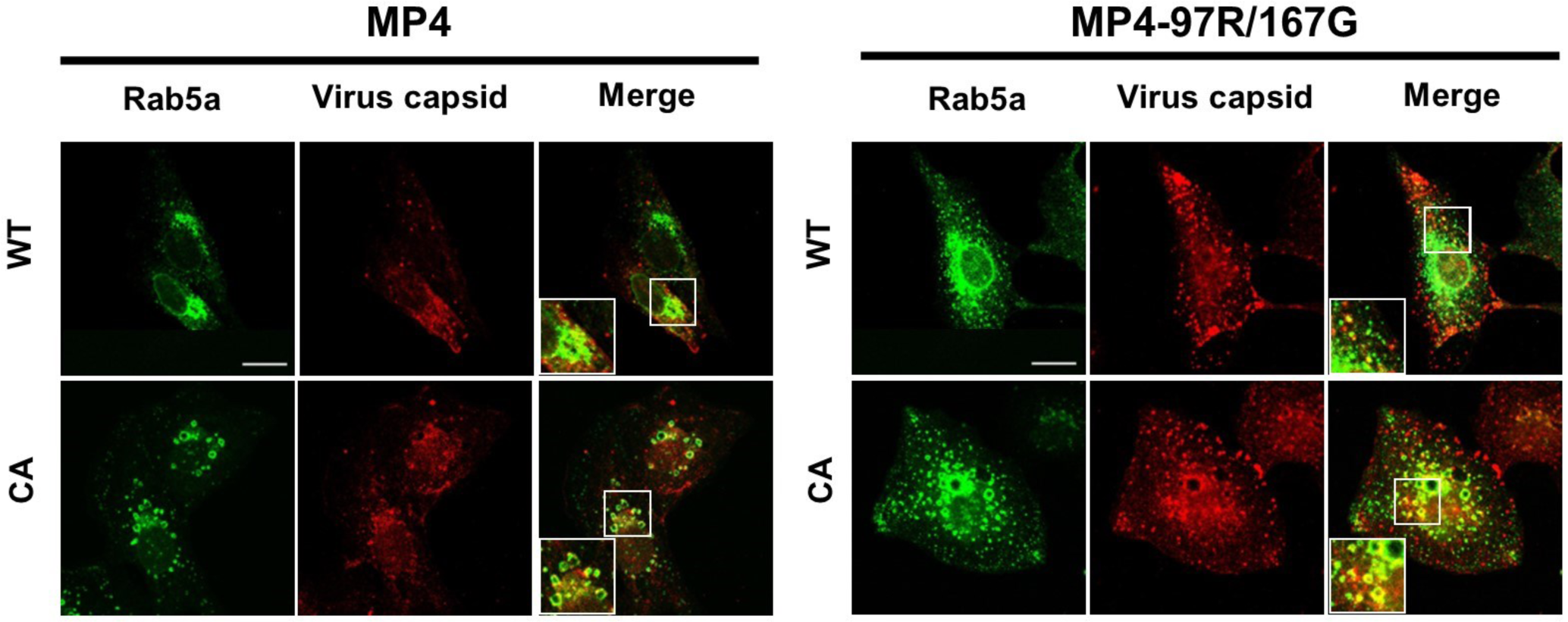
Viruses were detected in early endosomes at 30 mpi. Vero cells transiently expressing Rab5a-eGFP were infected with MP4 and MP4-97R/167G and fixed at 30 mpi. Colocalization of viruses was imaged with Rab5a in green and virus capsid (VP2) in red. Magnified area was highlighted in white box and displayed at left bottom of merged image.

**Fig S4.**
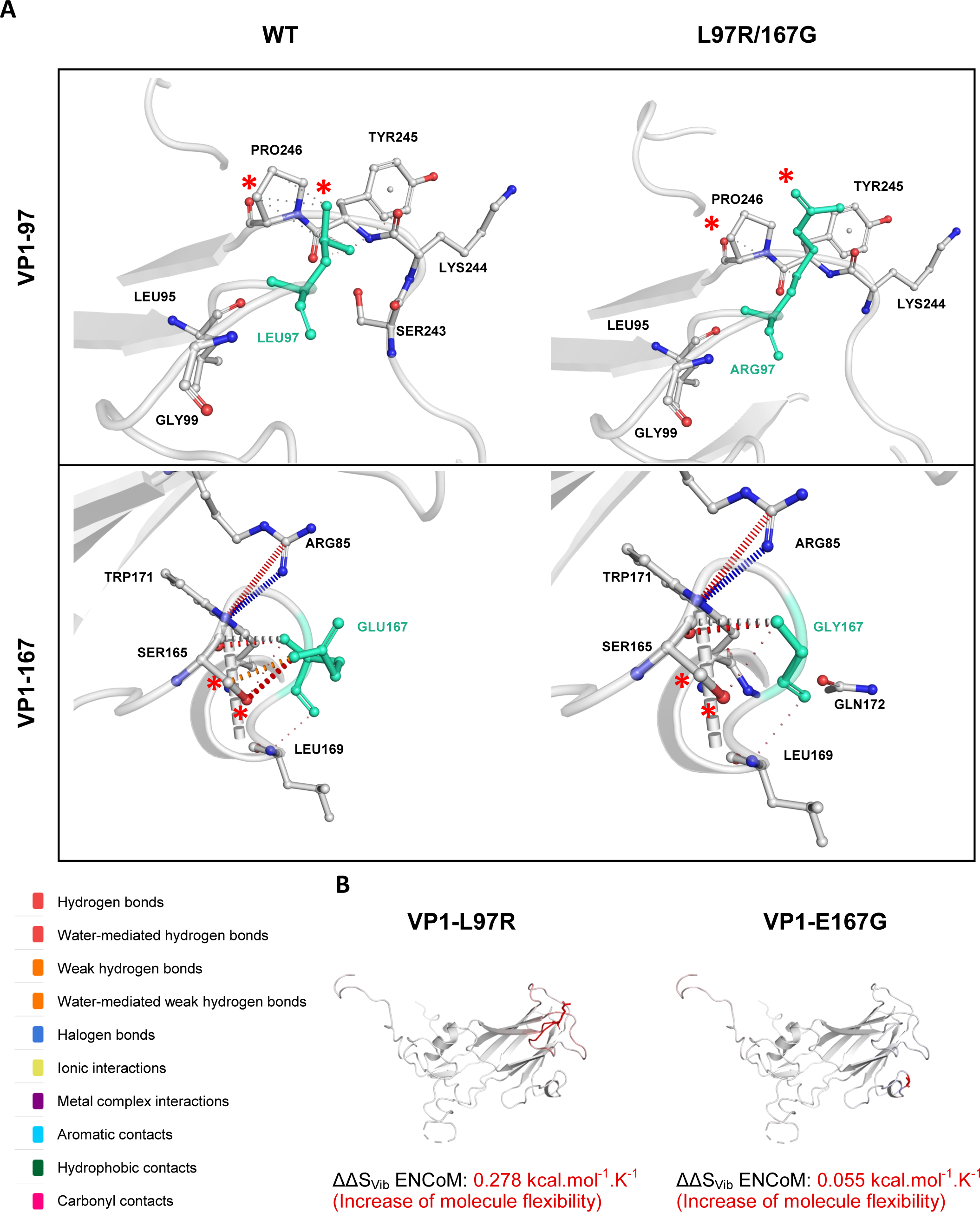
Visual presentation of DynaMut prediction of virus mutations on capsid amino acid interactions and protein stability. Prediction of changes in amino acid interactions and capsid stability induced by the VP1-L97R and VP1-E167G capsid mutations were performed using crystal structure of full assembled capsid on DynaMut server. (**A**) Interatomic interactions displayed and compared between WT and mutant capsid structures. VP1-97 and VP1-167 residues are labelled in light green and represented as sticks together with the surrounding interaction residues. Changes in interactions are highlighted on both WT and mutant structures with red asterisks (*) (**B**) VP1-L97R and VP1-E167G mutations decrease capsid stability. Computation of the vibrational entropy change (ΔΔS_Vib_) between WT and mutants. Amino acids in red indicate an increase of molecule flexibility.

**Table S1:**

| Mutation | pH | Temperature (°C) | Predicted free Gibbs energy change value ($\Delta\Delta G$) |
| --- | --- | --- | --- |
| VP1-L97R | 7 | 25 | -0.65<br>(Destabilizing) |
|  | 5 | 25 | -0.70<br>(Destabilizing) |
|  | 7 | 55 | -0.51<br>(Destabilizing) |
| VP1-E167G | 7 | 25 | -1.15<br>(Destabilizing) |
|  | 5 | 25 | -1.10<br>(Destabilizing) |
|  | 7 | 55 | -0.74<br>(Destabilizing) |
| VP1-E145Q | 7 | 25 | -0.80<br>(Destabilizing) |
|  | 5 | 25 | -0.89<br>(Destabilizing) |
|  | 7 | 55 | -0.71<br>(Destabilizing) |

